# Targeting EGFR in glioblastoma with a novel brain-penetrant small molecule EGFR-TKI

**DOI:** 10.1101/2021.01.09.426030

**Authors:** Jing Ni, Yanzhi Yang, Qiwei Wang, Johann S. Bergholz, Tao Jiang, Thomas M. Roberts, Nathanael S. Gray, Jean J. Zhao

## Abstract

Epidermal growth factor receptor (EGFR) is mutated or amplified in a majority of glioblastoma (GBM), and its mutation and focal amplification correlate with a more aggressive disease course. However, EGFR-directed tyrosine kinase inhibitors (TKIs) tested to date have yielded minimal clinical benefit. Here, we report a novel covalent-binding EGFR-TKI, CM93, as a potential drug to target adult GBMs with aberrant EGFR. CM93 has extraordinary brain exposure, with a brain-to-plasma ratio greater than 20-fold at estimated steady state. While all approved EGFR-TKIs are subject to extensive efflux transporter activity, CM93 does not inhibit the P-glycoprotein (P-gp) and breast cancer resistance protein (BCRP) efflux transporters in Caco-2 cells at expected clinically relevant plasma concentrations. Equally, CM93 demonstrates moderate absorption and permeation in Caco-2 cell monolayers with efflux ratios < 2, suggesting that it is not likely a substrate of an efflux transporter. Collectively, these *in vitro* data may account for the dramatic increase in brain exposure over plasma as noted above. Pre-clinical efficacy studies showed that CM93 is more effective than other EGFR-TKIs in blocking the proliferation of GBM tumor cells from both patient-derived and cultured human GBM cell lines with EGFR amplification and/or EGFRvIII mutation. In addition, CM93 administered as a single agent was able to attenuate the growth of orthotopic U251-EGFRvIII xenografts and extend the survival of tumor-bearing mice in a dose-dependent manner. Moreover, CM93 inhibited EGFR phosphorylation in GBM tumors derived from a novel genetically-engineered mouse (GEM) model of GBM with EGFRvIII expression both *in vitro* and *in vivo*. CM93 also extended the survival of mice bearing orthotopic allografts of GBM. Notably, mice maintained stable body weight during treatments with increasing doses of CM93 up to 75 mg/kg per day. Together, these data suggest that CM93 is a potential EGFR-TKI well suited for the treatment of adult GBM with mutant EGFR.

## Introduction

Glioblastoma (GBM) is the most common and malignant primary brain tumor in adults (Ostrom et al., 2019). Many targeted therapies have demonstrated extensive success in other cancer types but have limited efficacy in GBM, and the prognosis for patients with GBM remains grim (Kurz and Wen, 2018; Miller and Wen, 2016).

More than 50% of glioblastomas have aberrant EGFR genetic variants. Most of these EGFR variants occur through mutations in the extracellular domain (Vivanco et al., 2012). Among them, the most common EGFR variant (v), EGFRvIII (deletion of exon 2-7), has an in-frame extracellular domain truncation (Furnari et al., 2015). It has been shown that EGFR-mutant GBM cells are likely addicted to EGFR signaling (An et al., 2018; Huang et al., 2009). Therefore, EGFR is an attractive therapeutic target in GBM.

Currently, there are five EGFR tyrosine kinase inhibitors (TKIs; gefitinib, erlotinib, afatinib, dacomitinib and osimertinib) approved by the Food and Drug Administration (FDA) for the treatment of *EGFR*-mutant lung cancer in the United States. Gefitinib and erlotinib are first-generation EGFR-TKIs that inhibit catalytic activity by competing with ATP for binding to the ATP-binding site on the kinase domain, and which have significantly improved survival of patients over platinum chemotherapy. The second-generation of EGFR inhibitors, afatinib and dacomitinib, irreversibly inhibit all four ERBB receptors, including EGFR. As such, they are more potent inhibitors of EGFR but with increased toxicity (Le and Gerber, 2019). Osimertinib, the only FDA-approved third-generation EGFR TKI, is a covalent inhibitor designed to target EGFR resistance mutations that emerge with EGFR-TKI treatment (Le and Gerber, 2019).

While these first-and second-generation EGFR-TKIs have been shown to inhibit proliferation of GBM cells in preclinical experiments, they have not been effective in the clinic for GBM patients (Westphal et al., 2017). There are two principal reasons for these failures. The first reason involves important difficulties for drug penetration into the brain imposed by the blood-brain barrier (BBB). Osimertinib has been proposed for the treatment of EGFR-mutant GBMs in part because of its higher brain penetration (Wu et al., 2018), which likely contributes to its observed activity against brain metastases of lung cancer with EGFR mutations. However, dose-limiting toxicity (DLT) may prevent osimertinib from being a safe and effective drug for patients with GBM. Indeed, DLT highlights the second reason for the failure of these TKIs to effectively target GBM – namely, lack of a therapeutic window. Unlike EGFR mutations in lung cancers, such as exon-19 deletion, or L858R and T790M substitutions, which reside in the intracellular kinase domain (KD), the common feature of EGFR variants in GBM is a mutant extracellular domain with a wild type (WT) intracellular KD. Because of these complications, it has thus far been impossible to design a true targeted therapeutic that suppresses EGFR signaling within the tumor at concentrations that spare systemic WT EGFR function.

Here, we report CM93, a novel third-generation EGFR-TKI, that has favorable pharmacokinetic (PK) and safety profiles with extraordinary brain-specific distribution and accumulation (>2,000% brain penetration), which sets it apart from other reported EGFR inhibitors. Preclinical efficacy studies showed CM93 is more effective than other EGFR-TKIs at blocking the proliferation of GBM tumor cells from both patient-derived and cultured human GBM cell lines with EGFR amplification and/or EGFRvIII mutation. Our finding warrants further preclinical and clinical investigation for GBM.

## Results

### CM93 has high brain/plasma ratios

To determine the brain penetration of CM93, a pilot comparative assessment of brain exposure of CM93 and gefitinib was performed seven hours following a single oral dose of in rats with CM93 (30 mg/kg) or gefitinib (50 mg/kg). The results showed that CM93 brain/plasma (B/P) ratio was 28.3 and the B/P ratio for gefitinib was 0.22 **(Table 1)**. Given this encouraging finding, we proceeded with a more indepth comparative assessment of brain exposure of CM93 and gefitinib following continuous intravenous infusion to estimated steady state in rats. Both compounds were infused to a total dose of 3 mg/kg over five hours. Results from this study showed that CM93 B/P ratio was 20.3 and gefitinib was 0.55, consistent with our previous data from oral administration of the drugs **(Table 2)**. Notably, plasma concentrations of CM93 are much lower than that of gefitinib at all time points in both oral and intravenous administrations (**Tables 1 and 2**). These data confirm that CM93 is an ideal compound for treating GBM as it is preferentially present at high concentrations in brain with no apparent adverse effects in these pilot studies.

**Table 1.** Comparative Assessment of Brain Exposure of CM93 and Geftinib Following Single Oral (PO) Administration to Sprague Dawley Rats

| Test Article | Route | Dose (mg/kg) | Animal ID | Plasma Concentration by Hour (ng/mL) |  | Brain Tissue Concentration (ng/g) | Brain / Plasma (ratios) |
| --- | --- | --- | --- | --- | --- | --- | --- |
|  |  |  |  | 3 hr | 7 hr | 7 hr | 7 hr |
| Gefitinib | PO | 50 mg/kg Gefitinib | 1M001 | 1260 | 1030 | 562 | 0.546 |
|  |  |  | 1M002 | 2660 | 2240 | 561 | 0.250 |
|  |  |  | 1M003 | 1940 | 4700 | 666 | 0.142 |
|  |  |  | Mean | 1953 | 2657 | 596 | 0.224 |
|  |  |  | SD | 700 | 1870 | 60.3 | 0.0322 |
|  |  |  | n | 3 | 3 | 3 | 3 |
| CM93 |  | 30 mg/kg CM93 | 2M001 | 273 | 275 | 7400 | 26.9 |
|  |  |  | 2M002 | 181 | 265 | 8280 | 31.2 |
|  |  |  | 2M003 | 229 | 197 | 5210 | 26.4 |
|  |  |  | Mean | 228 | 246 | 6960 | 28.3 |
|  |  |  | SD | 46 | 42 | 1580 | 3.73 |
|  |  |  | n | 3 | 3 | 3 | 3 |

**Table 2.** Comparative Assessment of Brain Exposure of CM93 and Gefitinib Following Intravenous Administration to Sprague Dawley Rats

| Test Article | Route | Dose (mg/kg) | Animal ID | Plasma Concentration by Hour (ng/mL) |  |  |  |  | Brain Tissue Concentration (ng/g) | Brain / Plasma (ratios) |
| --- | --- | --- | --- | --- | --- | --- | --- | --- | --- | --- |
|  |  |  |  | 0.083 | 1 | 2 | 3 | 5 | 5 | 5 |
| Gefitinib | IV | 3 mg/kg | 1M001 | 39.9 | 135 | 210 | 279 | 292 | 151 | 0.517 |
|  |  |  | 1M002 | 38.2 | 125 | 166 | 206 | 282 | 149 | 0.528 |
|  |  |  | 1M003 | 35.8 | 88.4 | 161 | 216 | 303 | 189 | 0.624 |
|  |  |  | 1M004 | 55.6 | 115 | 161 | 225 | 286 | 155 | 0.542 |
|  |  |  | 1M005 | 42.9 | 122 | 164 | 265 | 281 | 153 | 0.544 |
|  |  |  | 1M006 | 39.8 | 116 | 180 | 252 | 237 | 129 | 0.544 |
|  |  |  | Mean | 42.0 | 117 | 174 | 241 | 280 | 154 | 0.550 |
|  |  |  | SD | 7.04 | 15.7 | 19.1 | 29.1 | 22.6 | 19.4 | 0.0377 |
|  |  |  | n | 6 | 6 | 6 | 6 | 6 | 6 | 6 |
| CM93 |  |  | 2M007 | 14.3 | 63.0 | 70.1 | 120 | 119 | 2640 | 22.2 |
|  |  |  | 2M008 | 14.2 | 63.5 | 93.4 | 95.5 | 107 | 2380 | 22.2 |
|  |  |  | 2M009 | 13.5 | 54.8 | 84.2 | 93.7 | 120 | 2400 | 20.0 |
|  |  |  | 2M010 | 12.8 | 59.6 | 81.6 | 106 | 109 | 2020 | 18.5 |
|  |  |  | 2M011 | 13.1 | 44.2 | 70.7 | 89.5 | 97.4 | 2000 | 20.5 |
|  |  |  | 2M012 | 11.3 | 51.1 | 71.9 | 94.7 | 109 | 1980 | 18.2 |
|  |  |  | Mean | 13.2 | 56.0 | 78.7 | 99.9 | 110 | 2240 | 20.3 |
|  |  |  | SD | 1.10 | 7.52 | 9.37 | 11.3 | 8.37 | 275 | 1.74 |
|  |  |  | n | 6 | 6 | 6 | 6 | 6 | 6 | 6 |

CM93 was a weak inhibitor of rosuvastatin transport via human breast cancer resistance protein (BCRP), with an apparent IC50 value of 3.02 μM. CM93 did not inhibit P-glycoprotein (P-gp)-mediated transport of digoxin (IC50 > 30.0 μM). CM93 also showed moderate permeability in Caco-2 cells. Based on these data, it is unlikely that CM93 is a substrate of efflux transporters, and there is generally a low risk of significant drug-drug interaction at therapeutic plasma concentrations via effects on these transporters.

### CM93 *in vitro* activity in HEK293 cells expressing EGFRvIII

As EGFRvIII is the most common EGFR variant in GBM, we set to test the effect of CM93 on the activity of EGFRvIII. We generated HEK293-EGFRvIII cells stably expressing EGFRvIII (293-EGFRvIII). CM93 reduced the phosphorylation of EGFRvIII at both tyrosine sites 1068 and 1173 (pEGFRvIII^Y1068^ and pEGFRvIII^Y1173^), as well as phosphorylation of the downstream signaling molecules ERK1 and ERK2 (ERK1/2) in a dose-dependent manner that was comparable to that of erlotinib **(Figure 1A)**. Further dose titration revealed an IC50 value of 0.19 μM for CM93 on EGFRvIII phosphorylation (**Figure 1B**). Furthermore, CM93 reduced viability of 293-EGFRvIII cells with an IC50 of 1.48 μM, which was lower than those of erlotinib (IC50 4.83 μM), gefitinib (IC50 15.67μM) and osimertinib (IC50 2.19 μM) (**Figure 1C**).

**Figure 1.**
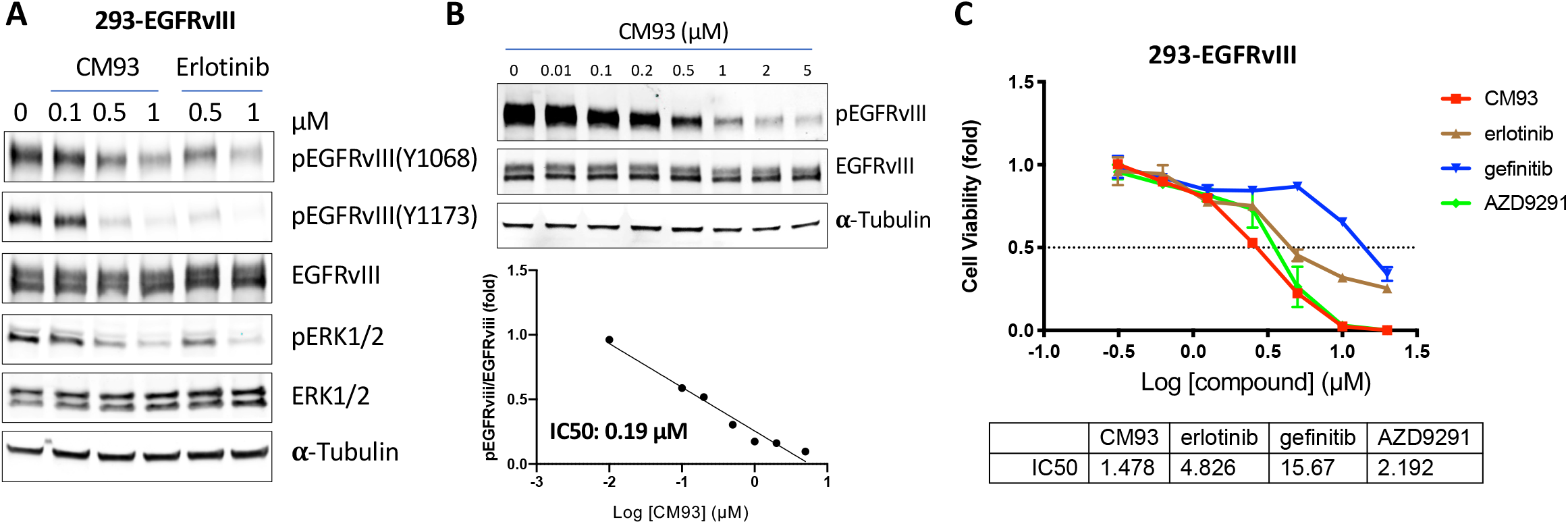
The effect of CM93 on 293 cells expressing EGFRvIII. (**A**) 293-EGFRvIII cells were treated with CM93 or erlotinib at the indicated doses for 6 hours and subjected to western blot analysis with the indicated antibodies. (**B**) 293-EGFRvIII cells were treated with CM93 at the indicated doses for 6 hours and subjected to western blot analysis with antibodies against pEGFRvIII^1068^ and EGFRvIII to determine IC50 for inhibition of pEGFRvIII^1068^. α-Tubulin was used as a loading control. (**C**) IC50 (µM) of CM93 and other EGFR-TKIs as indicated by the viability of 293-EGFRvIII cells.

### CM93 *in vitro* activity against GBM patient-derived cell lines harboring EGFR amplification and/or mutations

We next assessed CM93 on a panel of GBM patient-derived cell lines (PDCLs) harboring EGFR amplification and/or mutations with a neurosphere culture system. We extended this set of experiments to include lapatinib, as it was reported as a type II EGFR TKI that is highly active against GBM EGFR variants *in vitro* (Vivanco et al., 2012). Indeed, lapatinib more actively suppressed the survival of GBM patient-derived cells *in vitro* than the type I EGFR TKIs erlotinib and gefitinib. (**Figure 2A**). Notably, CM93 was the most potent TKI within this group, as shown by greater potency at reducing the viability of these PDCLs with the lowest IC50 values (**Figure 2B**)

**Figure 2.**
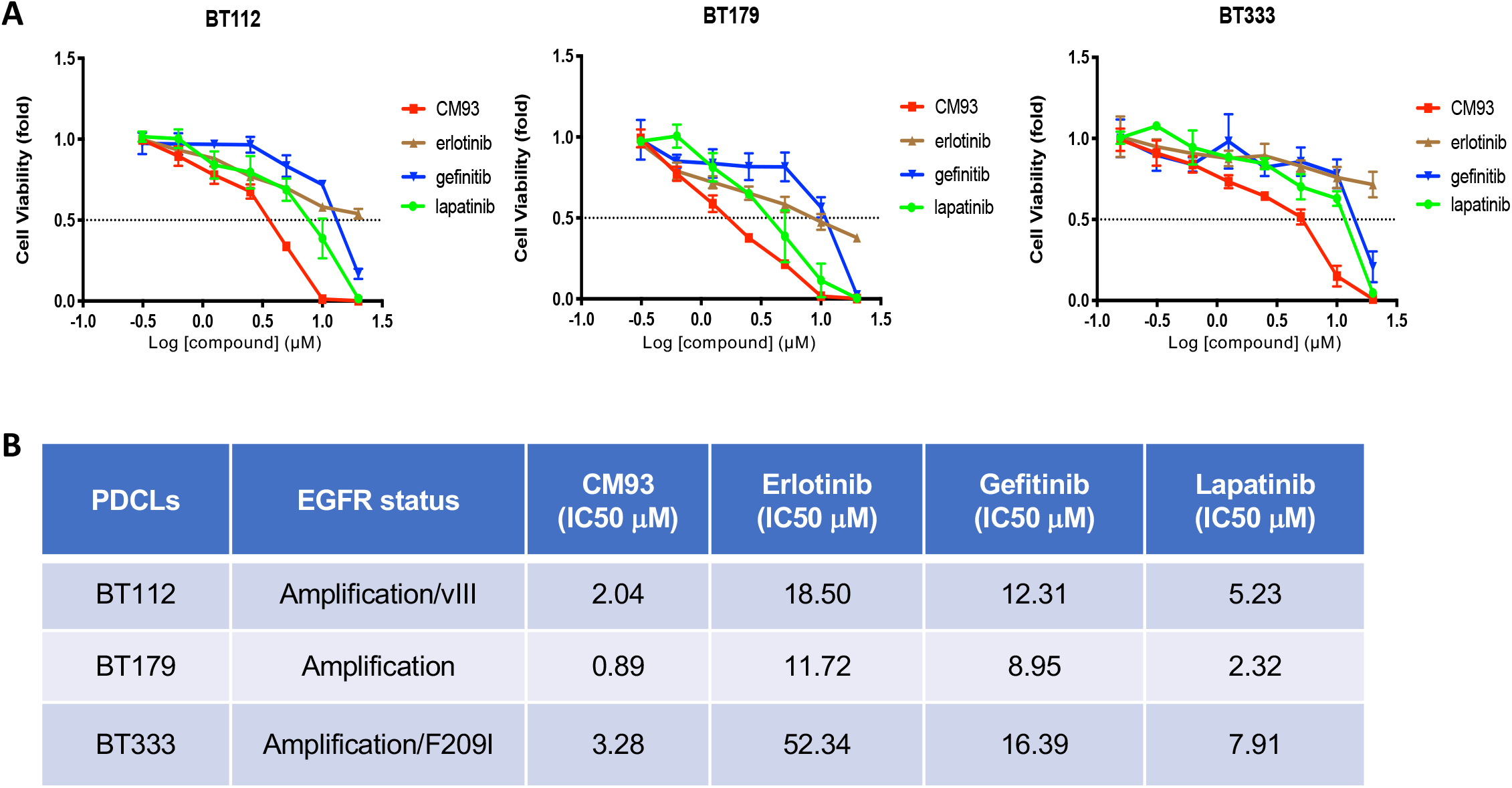
Effect of CM93 on cell viability in GBM PDCLs with EGFR amplification and/or mutation. (**A**) Viability of BT122, BT179 and BT333 cells upon treatment with CM93 or other EGFR inhibitors, as indicated, for 3 days in a dose titration from 0.156 μM to 20 μM for each drug. (**B**) IC50 of CM93 and other EGFR inhibitors for GBM PDCLs, BT112, BT179 and BT333 viability.

### CM93 *in vitro* activity against human GBM U251 cells expressing EGFRvIII

In parallel, we used the conventional human GBM U251 cell line as a surrogate model for both *in vitro* and *in vivo* studies. We engineered U251 cells to stably express EGFRvIII (U251-EGFRvIII) via retroviral-mediated gene transfer. CM93 was able to reduce EGFRvIII phosphorylation in U251-EGFRvIII cells in a dose-dependent manner in culture with an IC50 value of 0.174 μM. (**Figure 3A**). Examination of the effect of CM93 along with other EGFR-TKIs on the viability of U251-EGFRvIII cells revealed that, similar to the effect on 293-EGFRvIII cells (**Figure 1**), CM93 and osimertinib have comparable potencies on suppressing the viability of U251-EGFRvIII cells, with IC50 values of 1.52 μM and 2.64 μM, respectively, whereas erlotinib and gefitinib showed much higher IC50 values of 13.65 μM and 20.35 μM, respectively (**Figure 3B**)

**Figure 3.**
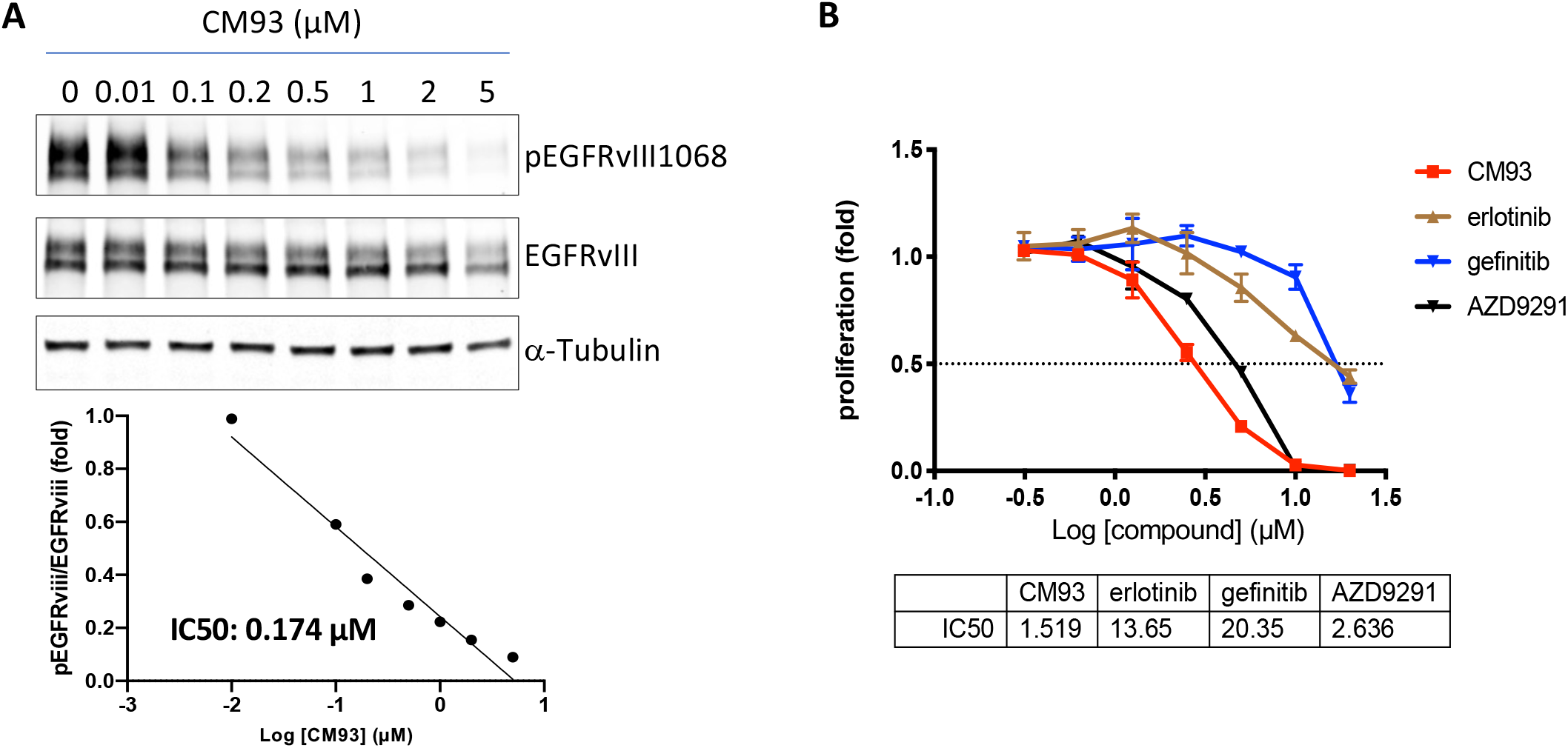
The effect of CM93 on GBM U251 cells expressing EGFRvIII *in vitro*. (**A**) U251-EGFRvIII cells were treated with CM93 at indicated doses for 20 hours and subjected to western blot analysis with the antibodies against pEGFRvIII^1068^ and EGFRvIII to determine IC50 for inhibition of pEGFRvIII^1068^. IC50 is 0.174 µM. (**B**) U251-EGFRvIII cells were treated with CM93 or other EGFR TKIs as indicated. IC50s of CM93 and other EGFR-TKIs are indicated on the table below the curves.

### CM93 *in vivo* activity against orthotopic xenograft model GBM U251-EGFRvIII

To evaluate the *in vivo* activity of CM93 in GBM with mutant EGFR, we generated U251-EGFRvIII GBM cells expressing luciferase to facilitate monitoring drug response *in vivo* by bioluminescence-imaging analysis. When luminescence signals were detectable, mice bearing orthotopic U251-EGFRvIII tumors were randomized into three treatment groups: vehicle control, CM93 at 37.5 mg/kg, or CM93 at 75 mg/kg. Two independent experiments were performed. Treatment with CM93 reduced the luminescence signals and prolonged the survival of the mice bearing orthotopic U251-EGFRvIII tumors in a dose-dependent manner in both cohorts (**Figure 4A-D**). The medium survival was 68 days for control mice, 80 days for mice treated with CM93 at 37.5 mg/kg, and 89.5 days for mice treated with CM93 at 75 mg/kg (**Figure 4E**). All mice appeared normal with no body weight loss during treatment time (**Figure 4F**), suggesting that CM93 has activity in orthotopic tumors of U251-EGFRvIII and is well-tolerated in mice.

**Figure 4.**
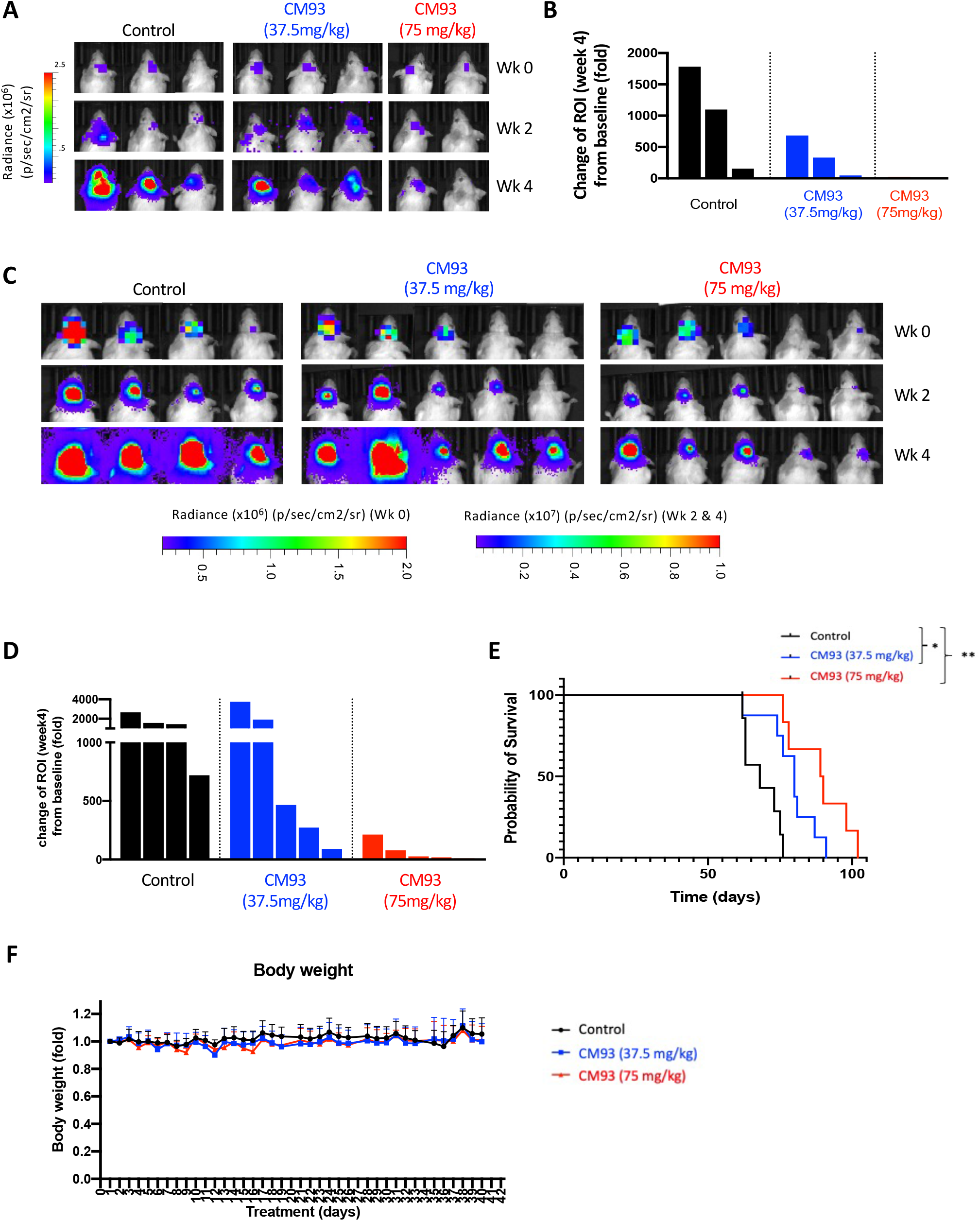
Effect of CM93 on orthotopic GBM xenograft model U251-EGFRvIII. (**A**) Representative bioluminescence imaging analysis of first cohort of mice bearing U251-EGFRvIII at indicated weeks after treatment with vehicle control (n=3), CM93 at 37.5 mg/kg (QD, n=3) or 75 mg/kg (QD, n=2). (**B**) Quantification of the regions of interest (ROI) in each mouse in the first cohort after treatment for four weeks compared to week zero, which was set as baseline. (**C**) Representative bioluminescence imaging analysis of second cohort of mice bearing U251-EGFRvIII at indicated times (weeks) after treatment with control (n=4), CM93 at 37.5 mg/kg (QD, n=5) or 75 mg/kg (QD, n=5). (**D**) Quantification of the regions of interest (ROI) in each mouse in the second cohort after treatment for four weeks compared to week zero, which was set as baseline. (**E-F**) Kaplan-Meier survival analysis (**E**) and body weight record (**F**) of U251-EGFRvIII xenograft-bearing mice from both cohorts treated with CM93 (37.5 mg/kg, PO QD, blue line, n=8), CM93 (75 mg/kg, PO QD, red line, n=7), or vehicle control (black line, n=7). Mean ± SD, ^*^ p < 0.05; ^**^ p < 0.01, Log-rank (Mantel-Cox) test.

### CM93 *in vitro* and *in vivo* activity against a genetically-engineered mouse (GEM) model of GBM

In GBM, EGFR mutations frequently coexist with CDKN2A deletion and PTEN deficiency (Brennan et al., 2013; Cancer Genome Atlas Research, 2008). We have generated a GEM model of GBM driven by *Cdkn2a* and *Pten* double deletion concomitant with *EGFRvIII* expression (termed CPEvIII). Notably, primary CPEvIII tumor cells can be cultured as neurospheres and also allografted intracranially in mice. As showed in **Figure 5A**, CM93 markedly reduced phosphorylation of EGFRvIII, ERK1/2 and S6RP in a dose-dependent manner *in vitro* in neurosphere cultures. CM93 (75 mg/kg, QD) also prolonged the survival of mice bearing intracranial orthotopic allograft of CPEvIII, with medium survival of 25.5 days for control mice and 33 days for CM93 treated mice, *p* = 0.017 (**Figure 5B**). Immunohistochemistry (IHC) analysis of pEGFR in tumors harvested from mice at the end point revealed that phosphorylation levels of EGFRvIII were reduced in mice treated with CM93 (**Figure. 4C**). CM93 was also well tolerated in this cohort of mice with no significant loss of body weight observed during CM93 treatment (**Figure 4D**). Together, these results suggest that CM93 has activity against CPEvIII GBM tumor cells both *in vitro* and *in vivo*.

**Figure 5.**
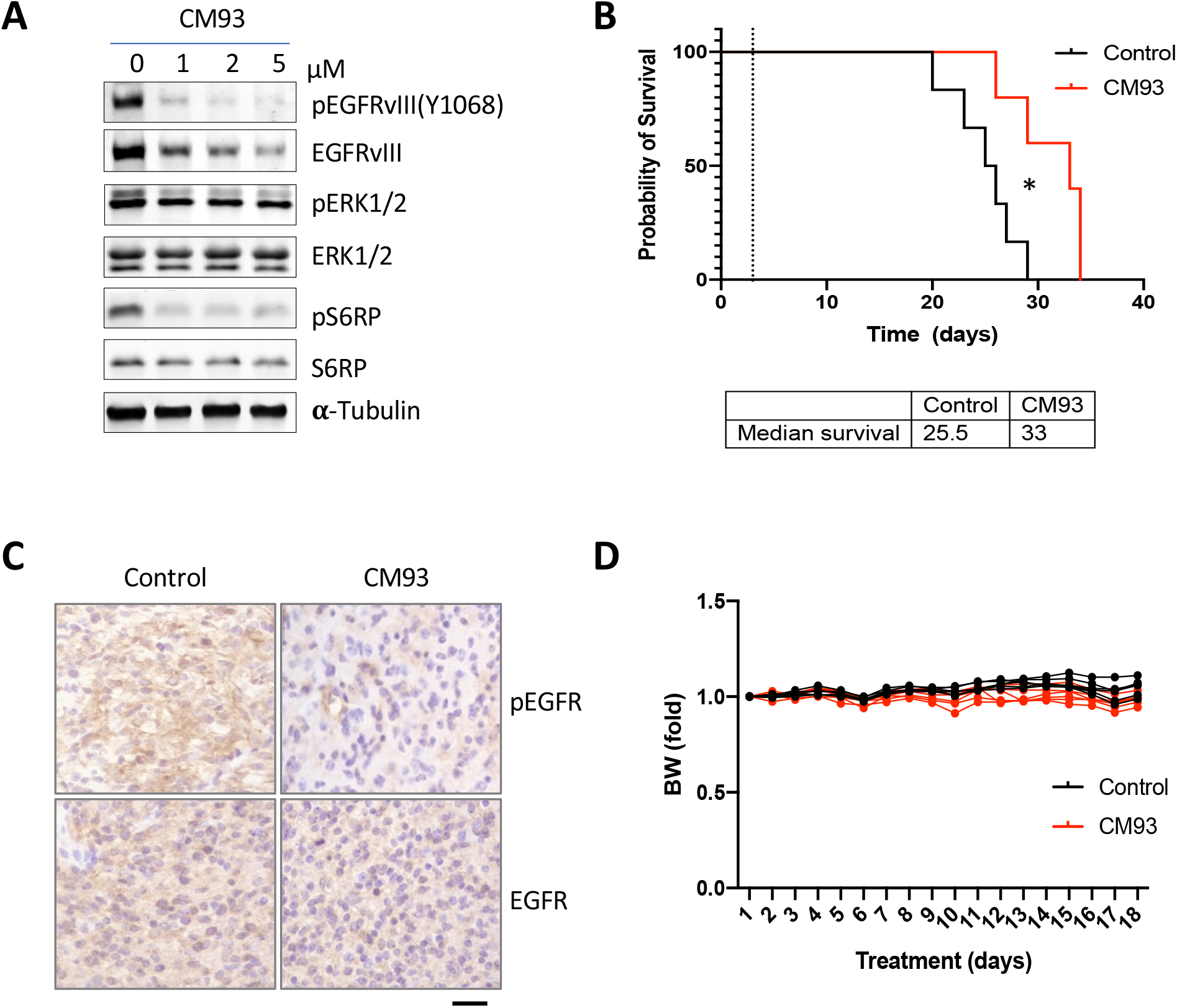
Effect of CM93 on a genetically-engineered mouse model of GBM driven by concurrent loss of *Cdkn2a* and *Pten* and expression of *EGFRvIII* (CPEvIII). (**A**) Primary mouse CPEvIII cells were treated with CM93 for 1 day. The cells were then subjected with western blot analysis. (**B**) Kaplan-Meier survival analysis of tumor-bearing mice treated with CM93 (75 mg/kg, PO QD, red line, n = 5) or vehicle control (black line, n = 6) \**p* < 0.05 (p = 0.017), Log-rank (Mantel-Cox) test. (**C**) IHC analysis of CPEvIII tumors collected at end point from mice treated with CM93 or vehicle control. Scale bar, 100 μm. (**D**) Body weight of tumor-bearing mice treated with CM93 or vehicle control.

## Discussion

Drug delivery across the BBB is a major barrier that all EGFR-targeting agents for GBM have to face. Numerous EGFR-TKIs have been evaluated for the treatment of GBM unsuccessfully, despite evidence that EGFR signaling is required for the viability of EGFR-mutant GBM cells (Westphal et al., 2017). Many of these inhibitors fail to cross the BBB or are substrates of drug efflux pumps, and often have a relatively small therapeutic window. Notably, CM93 distributes and accumulates in the brain at levels that are approximately 20-fold in excess of blood plasma levels. Unlike other EGFR-TKIs, which are subject to efflux transporters, CM93 is not a substrate of P-gp- or BCRP-mediated drug transport functions; instead, it is a modest inhibitor of BCRP. Moreover, CM93 exhibits a relatively high clearance rate to maintain relatively low plasma levels.

A recent study showed that, while EGFR is important for brain development during embryonic and early postnatal stages, adult mice with brain-specific deletion of EGFR appear to be normal (Robson et al., 2018), suggesting that EGFR inhibition in the CNS will not lead to dose-limiting toxicity. Thus, the distinct pharmacologic properties of CM93, in particular its high brain/plasma ratio, will potentially provide a “tissue-based” therapeutic window to allow effective inhibition of EGFR in the tumor, while relatively sparing the receptor systemically. Indeed, our pre-clinical data demonstrate that CM93 provides a sufficiently wide therapeutic window to effectively inhibit EGFRvIII intracranially without significant extracranial toxicity.

It has been well reported that the complexity of oncogenic signaling pathways, tumor heterogeneity and suppressive tumor immune microenvironment make GBMs refractory to most therapeutic agents (Buerki et al., 2018; Le Rhun et al., 2019). While CM93 as a single agent showed dose-dependent efficacy on orthotopic tumor models of GBM with EGFR/EGFRvIII, it provided a modest survival benefit to tumor-bearing mice. It is likely that CM93 in combination with drugs rational targeting other concurrent oncogenic pathways and/or immune modulatory agents will be required to improve the clinical outcome for GBM patients.

## Materials and methods

### Brain Exposure

Comparative assessment of brain exposure of CM93 and gefitinib following single Oral (PO) or intravenous (IV) administration to sprague dawley rats were performed by *Inotiv*. Protocols are available upon request.

### Cell culture

HEK293 cells and U251 cells were maintained in DMEM supplemented with 10% fetal bovine serum (FBS) and 100 μg/ml penicillin-streptomycin. Mouse neural stem cells (NSCs) were expanded in NeuroCult proliferation medium (mouse) (StemCell Technologies) supplemented with 20 ng/ml EGF. Primary mouse glioma cells (*CPEvIII*) were cultured in NeuroCult proliferation medium (mouse) (StemCell Technologies) supplemented with 20 ng/ml EGF, 10 ng/ml FGF and 0.0002% Heparin. Primary human glioblastoma lines BT112, BT179 and BT333 were maintained in NeuroCult proliferation medium (human) (StemCell Technologies) with 20 ng/ml EGF, 10 ng/ml FGF and 0.0002% Heparin.

### Compounds and Reagents

Gefitinib and erlotinib were purchased from Selleck Chemicals. Lapatinib was purchased from MedChemexpress. Osimertinib (AZD9291) was obtained from commercial sources (Wang et al., 2020). CM93 was synthesized by Pharmaron (Wang et al., 2020). For *in vitro* studies, compounds were dissolved in DMSO. For *in vivo* studies, CM93 was dissolved in 10% NMP / 90% PEG300 and administered by oral gavage at 37.5 mg/kg or 75 mg/kg daily.

### Western blot analysis

Western blot analysis was performed as described previously (Ni et al., 2012; Ni et al., 2016; Ni et al., 2017). Anti-pEGFR-1068 (#3777), anti-pEGFR-1173 (#4407), anti-EGFR (#4267), anti-pERK1/2 (#9101), anti-ERK1/2 (#9102), anti-S6RP (#2211), and anti-S6RP (#2217) antibodies were purchased from Cell Signaling Technology. Anti-α-Tubulin antibody was purchased from Sigma.

### Cell viability assay

Cells were seeded in 96-well plates at a density of 1,000 cells per well and treated with two-fold serial dilutions of compounds with a starting concentration of 20 μM. Cell viability was assessed after three days of treatment by CellTiter-Glo (Promega). Curve fitting analysis and IC_50_ value determination were performed using GraphPad Prism 8.

### Mice

*Pten*^*f/f*^ (from Dr. Hong Wu, UCLA) mice were backcrossed to C57BL/6 strain background for 10 generations. They were then crossed with *Cdkn2a-null* (*Ink4a*^*-/-*^*;/Arf*^*-/-*^) mice (Ni et al., 2017), which are on an C57BL/6 background to produce *Cdkn2a-null;Pten*^*f/f*^ mice. ICR-SCID mice were purchased from Taconic. All animal experiments were performed in accordance with NIH animal use guidelines and protocols approved by the Dana-Farber Cancer Institute Animal Care and Use Committee (IACUC).

### Intracranial injections of cells

Cells (100,000 cells resuspended in 1 μl PBS) were intracranially injected into the right striatum (0 mm anterior, 2 mm lateral, and 2.5 mm ventral to bregma) of 8-10 week-old ICR-SCID mice. Animals were monitored daily for development of neurological defects.

### Primary mouse CPEvIII glioma

Neural stem cells (NSCs) from E14.5 embryonic mice (*Cdkn2a-null;Pten*^*f/f*^) striata were isolated and cultured as previously described (Rietze and Reynolds, 2006). NSCs were infected twice with adenovirus expressing Cre recombinase (AdCre; MOI50) (University of Iowa) to knock out floxed *Pten*. Cells were then transduced with retrovirus expressing EGFRvIII (pBabe-puro-EGFRvIII) (from Dr. Charles Stiles, DFCI) and selected with 1 μg/ml puromycin. The resulting cells (*Cdkn2a*^*null*^; *Pten*^*null*^; *EGFRvIII*, denominated CPEvIII) can form a glioma after grafted into the mouse brain. Tumors were then isolated and mechanically dissociated for expansion *in vitro* and *in vivo*.

### Bioluminescence imaging

Cells were transduced with lentiviral luciferase (HIV-Luc-zsGreen, addgene#39196). Bioluminescence signals from luciferase-expressing cells in live mice were recorded 10 minutes after intraperitoneal injection of D-luciferin (80 mg/kg) (Gold Biotechnology) with IVIS Lumina III Imaging System (PerkinElmer). The signals were analyzed with Living Image Software (PerkinElmer).

### Statistical Analysis

Statistical analysis of animal survival was determined by the log-rank (Mantel-Cox) test (Prism). Data were considered statistically significant when *P* < 0.05.

## Acknowledgements

We thank Drs Jose McFaline-Figueroa, Charles Stiles and Patrick Wen at Dana-Farber Cancer Institute and Drs. William D. Kerns and Xiang Y. Yu at Accellient for scientific discussions. We thank Dr Keith Ligon for providing patient-derived cell lines.

